# Piwi mutant germ cells transmit a form of heritable stress that promotes longevity

**DOI:** 10.1101/326769

**Authors:** Bree Heestand, Ben McCarthy, Matt Simon, Evan H. Lister-Shimauchi, Stephen Frenk, Shawn Ahmed

## Abstract

The *C. elegans* Argonaute protein PRG-1/Piwi and associated piRNAs protect metazoan genomes by silencing transposons and other types of foreign DNA. As *prg-1* mutants are propagated, their fertility deteriorates prior to the onset of a reproductive arrest phenotype that resembles a starvation-induced stress response. We found that late-generation *prg-1* mutants with substantially reduced fertility were long-lived, whereas early- or mid-generation *prg-1* mutants had normal lifespans. Loss of the stress response transcription factor DAF-16 caused mid- or late-generation *prg-1* mutants to live very short lives, whereas overexpression of DAF-16 enabled both mid- and late-generation *prg-1* mutants to live long. Cytoplasmic P-bodies that respond to stress increased in long-lived late-generation *prg-1* mutants and were transmitted to F1 but not F2 cross-progeny. Moreover, moderate levels of heritable stress shorten late-generation *prg-1* mutant longevity when DAF-16 or P bodies are deficient. Together, these results suggest that the longevity of late-generation *prg-1* mutants is a hormetic stress response. However, dauer larvae that occur in response to stress were not observed in late-generation *prg-1* mutants. Small germ cell nucleoli that depended on germline DAF-16 were present in late-generation *prg-1* mutants but were not necessary for their longevity. We propose that *prg-1* mutant germ cells transmit a form of heritable stress, high levels of which promote longevity and strongly reduce fertility. The heritable stress transmitted by PRG-1/Piwi mutant germ cells may be generally relevant to epigenetic inheritance of longevity.

**Core message of paper:** *prg-1*/Piwi mutants with strongly reduced fertility live long and longevity is transmitted for one generation to F1 cross progeny. Stress granules are increased and germ cell nucleoli are small for long-lived Piwi mutants and their F1 progeny. Loss of *daf-16* stress response transcription factor or *dcap-1* P body protein causes very short life for worms when *prg-1* mutant fertility is moderately reduced, whereas moderate fertility is sufficient to extend lifespan when somatic DAF-16 is overexpressed. We propose that *prg-1* mutant germ cells transmit a heritable epigenetic factor that is stressful and elicits two hormetic stress responses: reproductive arrest and longevity.

## Introduction

In principle, aging can be divided into two categories: aging of post-mitotic cells, like neurons or muscle cells, and aging of cells that proliferate, like hematopoietic stem cells. Proliferative aging occurs for human fibroblasts that display a limited number of cell divisions when grown *in vitro* (40-60), termed the Hayflick limit, which occurs as a consequence of telomere erosion (Shay & Wright 2019). A few short telomeres that arise in proliferating human fibroblasts are sufficient to trigger senescence, which is a state of permanent cell cycle arrest that suppresses tumor development (Campisi 2011). However, senescence can also be induced by other stresses, including aberrant cellular proliferation conditions and non-telomeric forms of DNA damage (Micco et al. 2021). Non-telomeric forms of stress transmitted by mammalian germ cells might influence the proliferative aging of somatic cells.

The relationship between proliferative and post-mitotic aging is not well understood. Germ cells are an effectively immortal cell lineage that can be transmitted indefinitely. The fertility of *C. elegans mortal germline* (*mrt*) mutants declines over a number of generations and ultimately results in complete sterility. Some *mrt* mutants are deficient for telomerase-mediated telomere maintenance and their germ cells transmit chromosome termini that shorten progressively, ultimately eliciting end-to-end chromosome fusions and genome catastrophe (Meier et al. 2006). We previously examined post-mitotic aging of *C. elegans* adults that were deficient for the telomerase reverse transcriptase, *trt-1,* and found that early-, middle- and late-generation *trt-1* mutants had wild-type lifespans, even though the fertility of late-generation *trt-1* mutants was severely compromised (Meier et al. 2006). Therefore, telomere erosion triggers proliferative aging and senescence of mammal somatic cells but does not impact aging of post-mitotic cells in *C. elegans* adults.

A second germ cell immortality pathway is regulated by the *C. elegans* Argonaute protein Piwi/PRG-1, which associates with piRNAs to promote genomic silencing in germ cell nuclei (Simon et al. 2014). Piwi/PRG-1 localizes to perinuclear P granules of germ cells and interacts with thousands of 21 nucleotide piRNAs that scan the genome for foreign nucleic acids such as transposons, viruses and transgenes (Reed et al. 2020). PRG-1 and associated piRNAs can detect genomic intruders based on small genomic incongruities that occur when homologous chromosomes are paired during meiosis (Leopold et al. 2015). mRNAs that are targeted by piRNAs are utilized to promote biogenesis of 22 nucleotide effector siRNAs that typically possess a 5’ guanine (22G RNAs), which promote genomic silencing via nuclear RNAi factors and histone modifying proteins (Billi 2014).

Freshly outcrossed *prg-1*/Piwi mutants display normal fertility for a number of generations, but their fertility then progressively declines and ultimately culminates in complete sterility (Simon et al. 2014; Barucci et al. 2020; Reed et al. 2020). The PRG-1/Piwi pathway promotes silencing of foreign genetic elements like transposons whose expression might promote *prg-1* mutant sterility (Simon et al. 2014; McMurchy et al. 2017). However, PRG-1/piRNAs also repress the inappropriate silencing of some germline genes, possibly in response to an imbalance of silencing factors and 22G RNAs that arises in the absence of Piwi/piRNAs (Montgomery et al. 2021; de Albuquerque et al. 2015; Barucci et al. 2020; Reed et al. 2020). Moreover, sterility occurs immediately if fertile *prg-1* mutants that lack all 22G RNAs are crossed to restore 22G RNA production, suggesting that excessive 22G RNA levels cause *prg-1* mutant sterility (de Albuquerque et al. 2015). Although 22G RNAs antisense to histone and ribosomal RNA accumulate in late-generation *prg-1* mutants (Barucci et al. 2020; Wahba et al. 2021), histone or ribosomal RNA levels may not cause the sterility of late-generation *prg-1* mutants (Montgomery et al. 2021). Therefore, the precise cause of *prg-1* mutant sterility is not understood but may be due to inappropriately high levels of one or more 22G RNA classes.

The fertility of a small fraction of sterile late-generation *prg-1* mutants can be revived if the sterile adults are placed on a different food source, which indicates that *prg-1* mutant sterility is a form of reproductive arrest (Heestand et al. 2018). Environmental stresses like starvation or seasonal cues can trigger a state of adult reproductive arrest termed Adult Reproductive Diapause (Angelo & Van Gilst 2009). Rapid shrinking of the germ line occurs for sterile *prg-1* mutants at the L4 to adult ransition (Heestand et al. 2018), and this is also observed for wild-type *C. elegans* worms that are starved as L3 or L4 larvae and mature into sterile adults whose fertility can be revived by addition of food (Angelo & Van Gilst 2009). The reproductive arrest phenotype of late-generation *prg-1*/Piwi mutants might therefore occur in response to a form of heritable endogenous stress that is transmitted by Piwi mutant germ cells (Heestand et al. 2018).

Wild type *C. elegans* worms subjected to dietary restriction, acute starvation or other stresses can live longer than unstressed animals (Baugh & Hu 2020). These observations are consistent with the concept of hormesis, where longevity is extended by biological responses to non-debilitating levels of stress or toxins (Shore & Ruvkun 2013; Calabrese et al. 2015; Masoro 2007). Here we show that *prg-1*/Piwi mutant germ cells transmit a form of heritable stress that promotes adult longevity.

## Results

### Late-generation *prg-1* mutants are long-lived

The transgenerational decline of *prg-1*/Piwi mutant fertility is stochastic (Simon et al. 2014). If one propagates thirty *prg-1* mutant strains derived from a single outcrossed *prg-1* mutant F2 animal, then at generation F20 some will have normal brood sizes, some will have medium brood sizes will live normal lifespan and some will have small brood sizes and be long-lived. Therefore, the precise generation number of animals used in assays in this paper is arbitrary and misleading, because it is the level of fertility of *prg-1* mutant strains, rather than the number of generations that they have grown, that dictates whether they will live normal or long lives. We assess *prg-1* mutant fertility by placing 4 or 6 L1 larvae on an NGM plate seeded with *E. coli* and qualitatively scoring worm abundance after a week of growth at 20°C. ‘Early-generation’ strains have normal levels of fertility and are close to starvation after one week. ‘Mid-generation’ strains display modest decreases in fertility that result in some *E. coli* food remaining after a week. ‘Late-generation’ *prg-1* mutant strains display a pronounced reduction in fertility, frequent males, and plentiful *E. coli* food left after a week, whereas sterile *prg-1* mutants are dark adults that lack embryos and have an empty transparent uterus that appears white on a dissecting microscope (Figure 1A, described further in Supplemental Experimental Procedures).

**Figure 1.**
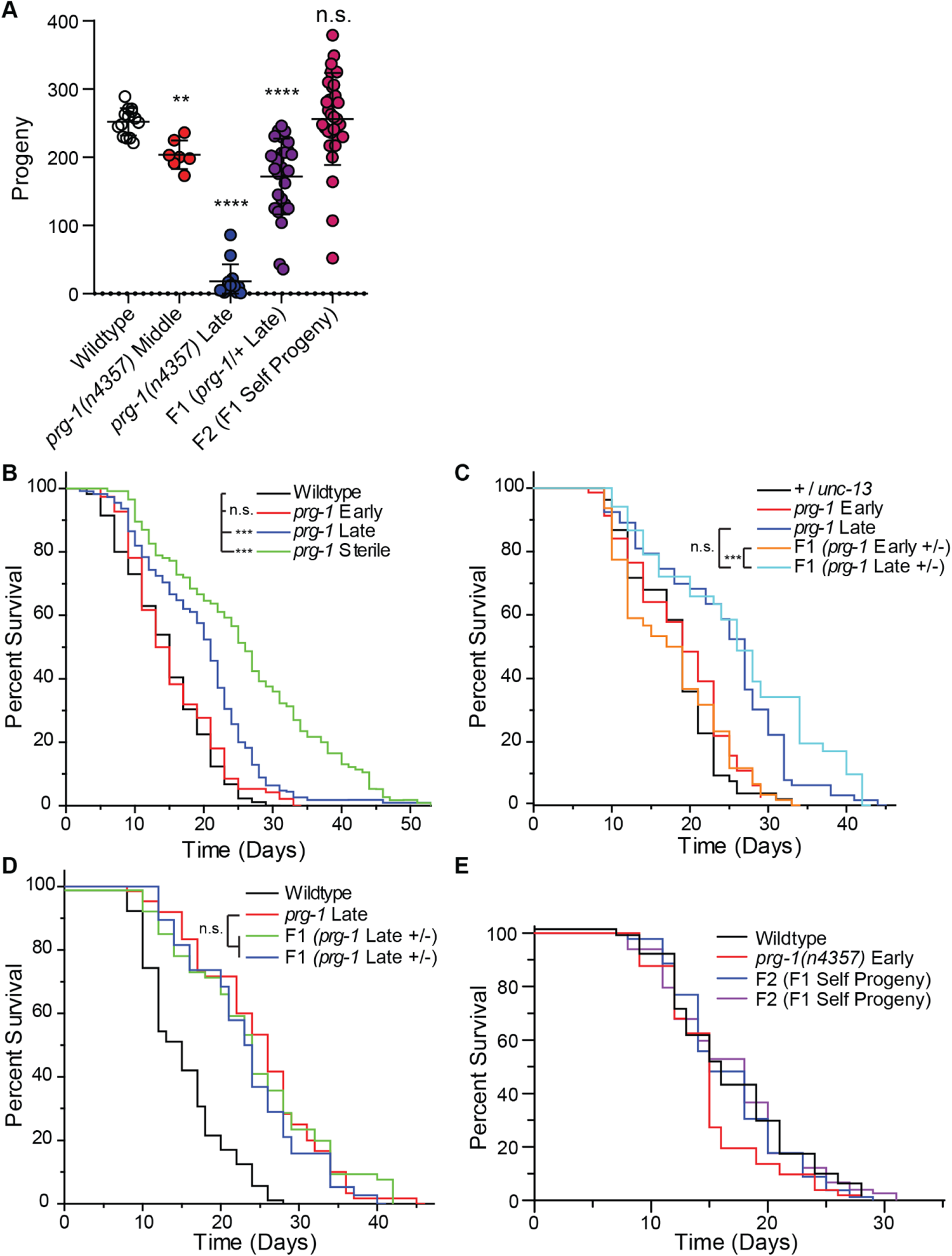
*prg-1* mutants show a transgenerational increase and epigenetic inheritance of lifespan. (A) Brood size counts for wildtype (N2), *prg-1(n4357)* mutant middle generation, *prg-1(n4357)* late generation, F1 progeny from late generation *prg-1* mutants crossed to wildtype males, and F2 self progeny from F1 mothers. **p=0.0024, ****p<0.0001, n.s. is not significant. Lifespan curves for (B) early, late, and sterile generation *prg-1* mutants, (C) +/*unc-13*, early or late generation *prg-1*, and early or late *prg-1* +/-F1 progeny, (D) two additional independent lines of heterozygous (*prg-1* +/-) late generation F1 progeny compared to homozygous non-crossed sisters (p=1), and (E) F2 self progeny from late *prg-1* +/-F1 (p=0.2069). n.s. is not significant, *** p<0.001, p values determined using Log-rank (Mantel-Cox) test and reported in S1 Table.

When initiating a late-generation *prg-1* mutant lifespan assay, we would examine thirty F18 plates after a week of growth and select those plates with small brood sizes and plentiful *OP50 E. coli* food. We would transfer four F19 L4 larvae from each small brood size plate to fresh plates seeded with *OP50*, F20 brood size would be scored to ensure that it remains small, and lifespan assays would be initiated with F20 L4 hermaphrodite larvae. Although sterile animals arise in late-generation *prg-1* mutant strains with small brood sizes, we studied the lifespan of fertile late-generation *prg-1* mutants by selecting against sterile adults lacking embryos.

We hypothesized that *C. elegans prg-1* mutants become sterile in response to a heritable stress that might affect aging. We found that longevity of early-generation *prg-1* mutant adults was not significantly different in comparison to wildtype controls (Figure 1B, S1 Table, p=1), whereas fertile and sterile late-generation *prg-1*/Piwi mutant adults displayed significant 46.5% and 81.8% increases in median lifespan, respectively (Figure 1B, S1 Table, p=0.0008 and p=1.55E-16). In contrast, middle-generation *prg-1* mutants that have modestly decreased fertility (Figure 1A) did not live longer than wildtype controls (Figure 2B, p=0.3708).

**Figure 2.**
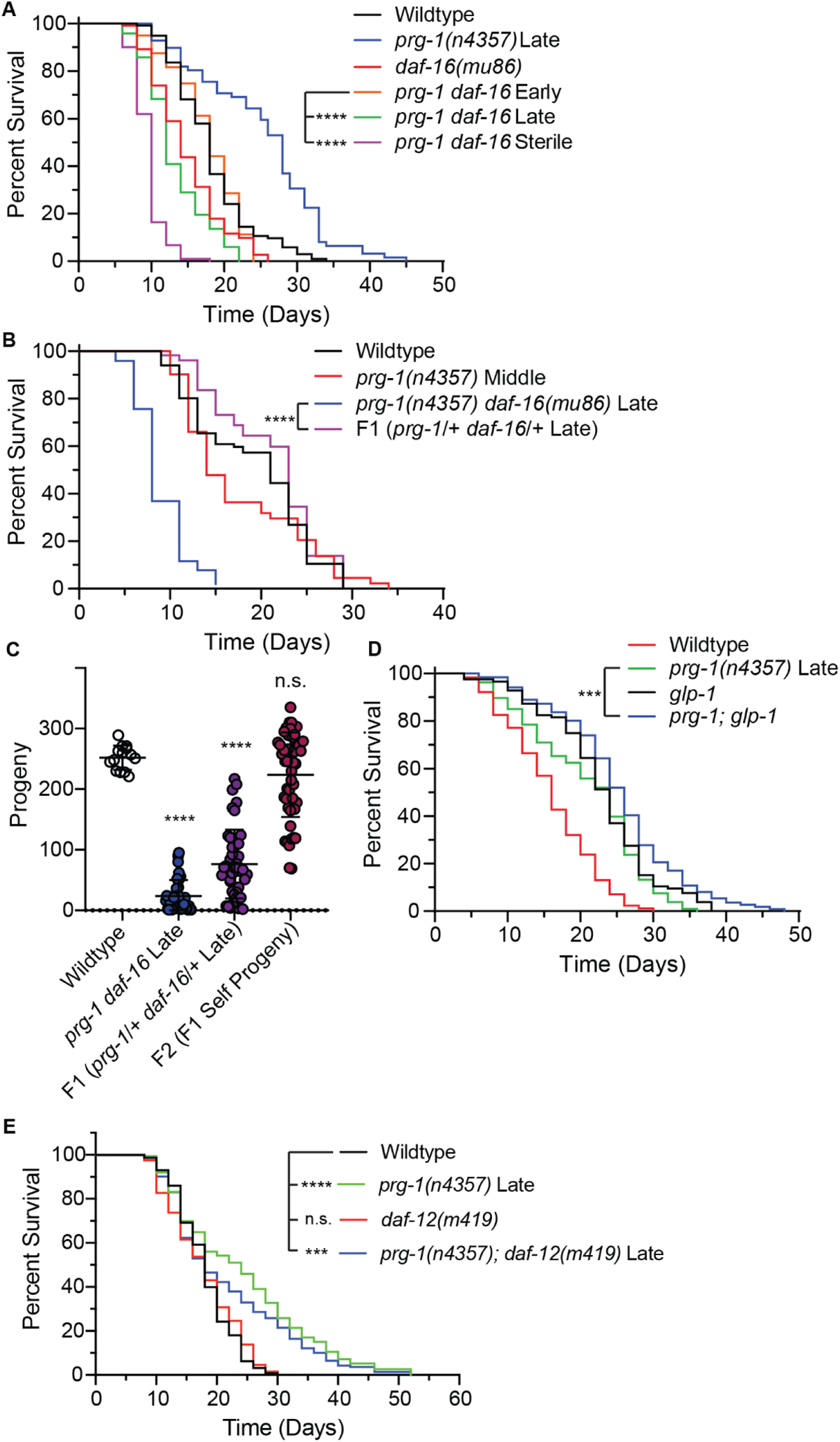
DAF-16 is required for heritable stress induced longevity. (A) Lifespan curves for wildtype, late generation *prg-1(n4357)*, *daf-16 (mu86)*, and early-late- and sterile generation *prg-1(n4357) daf-16(mu86*), (B) *prg-1(n4357)* middle generation (p=n.s. vs WT), late generation *prg-1(n4357) daf-16(mu86)*, and the F1 progeny of late generation *prg-1(n4357) daf-16(mu86)* crossed with WT males. (C) Brood size counts for wildtype (N2), late generation *prg-1(n4357) daf-16(mu86)*, F1 worms from late generation *prg-1 daf-16* mutants crossed to wildtype males, and F2 self progeny from F1 mothers. (D) Lifespan curve for wildtype, late generation *prg-1(n4357)*, *glp-1(e2141)*, and *prg-1(n4357); glp-1(e2141)* double mutants. (E) Lifespan curve for wildtype, *daf-12(m419)*, late generation *prg-1(n4357)*, and late generation *prg-1(n4357); daf-12(m419)* double mutants. ****p<0.0001, ***p<0.001, *p<0.05, n.s. is not significant.

### *prg-1* mutant germ cells transmit longevity to F1 cross-progeny

We asked if *prg-1* mutant germ cells transmit a factor that affects longevity by crossing three independently derived lines of late-generation *prg-1* mutant hermaphrodites with wild-type males that were heterozygous for an *unc-13* mutation, which is tightly linked to *prg-1*. To ensure that the presence of the *unc-13* balancer mutation did not confer a dominant effect on longevity, we also measured the lifespan of *unc-13 +/-* heterozygous controls and found that their longevity was not significantly different from that of early-generation *prg-1* mutant controls (Figure 1C, S1 Table, p=1). All three independent lines of late-generation *prg-1* mutants gave rise to *prg-1* +/-F1 progeny with significant increases in lifespan in comparison with *unc-13 +/-* controls, with early-generation *prg-1* mutants, as well as with early-generation *prg-1* +/-F1 controls (Figure 1C,D, S1 Table, p<0.05 in all comparisons). The lifespan of late-generation *prg-1* +/-F1 cross progeny was not different than that of their late-generation *prg-1 -/-* mutant parents (Figure 1C,D, S1 Table, p>0.72 in all cases). Late-generation *prg-1* mutant adults give rise to few progeny, but their *prg-1* +/-F1 cross progeny had a significant increase in brood size, comparable to middle-generation *prg-1* mutant strains that were not long-lived (Figure 1A).

We studied the F2 progeny of late-generation *prg-1 - / -* mutants using a tightly linked *unc-13* mutation to select against homozygous wild type *prg-1 + / +* animals in the F2 generation. We found that brood size of a mixture of heterozygous *prg-1 + / -* and homozygous *prg-1 - / -* F2 adults was equivalent to wildtype (Figure 1A) and that their lifespan was the same as that of wildtype controls, even though ⅓ of these animals should have been *prg-1 - / -* mutant homozygotes (Figure 1E). Therefore, the epigenetic defect of late-generation *prg-1* mutants that induces longevity is transmitted to their F1 cross-progeny but is reset by the F2 generation. This demonstrates that *prg-1* is not haploinsufficient for fertility or longevity phenotypes observed in the F1 progeny of late-generation *prg-1* mutants. Rather, an epigenetic factor present in late-generation *prg-1*/Piwi mutant germ cells is transmitted for a single generation to promote longevity.

We tried asking if late-generation *prg-1* mutant males could transmit longevity to F1 cross progeny but found that late-generation *prg-1* mutant males would not create cross-progeny, even when mated with hermaphrodites that are mutant for *unc-3*, which mates readily with N2 wild-type males. Therefore, we were unable to determine if the longevity transmitted by *prg-1* mutants to their F1 progeny is only transmitted maternally or if it is a form of nuclear inheritance that is transmitted by both male and female gametes.

### Heritable stress is detrimental to longevity in the absence of *daf-16*

The DAF-16 transcription factor that promotes longevity and stress resistance (Baugh & Hu 2020) interacts in three distinct ways with *prg-1/*Piwi. First, starvation of *prg-1* mutants for one day per week from the F4 generation onwards suppresses the transgenerational sterility of *prg-1* mutants by activating DAF-16 (Simon et al. 2014). However, activation of DAF-16 in late-generation *prg-1* mutants does not suppress their sterility (Heestand et al. 2018). Second, sterile *prg-1* mutants display a pronounced germ cell atrophy phenotype as L4 larvae mature into 1 day old adults, but deficiency for *daf-16* significantly reduces levels of germ cell atrophy (Heestand et al. 2018). A third function of DAF-16 occurs for 5 day old sterile *prg-1* mutants, which are in a state of prolonged reproductive arrest. A small fraction of day 5 sterile adults can become fertile if moved to an alternative food source, which requires DAF-16 (Heestand et al. 2018).

Given that DAF-16 promotes longevity in several contexts (Baugh & Hu 2020), we asked if DAF-16 promotes longevity of late-generation *prg-1* mutants. We found that late-generation and sterile generation *prg-1 daf-16* double mutant animals lived significantly shorter than either early-generation *prg-1 daf-16* double mutants (late p=3.53E-12 and sterile p=1.55E-35) or *daf-16* single mutant controls (late p=0.028 and sterile p=2.71E-19), (Figure 2A, S1 Table). However, when late-generation *prg-1 daf-16* double hermaphrodites were crossed with wild-type males, their F1 cross-progeny were longer lived than their parents (Figure 2B, p<0.0001). Like progeny of late-generation *prg-1* mutants, the brood size of *prg-1 daf-16* cross progeny was significantly reduced for F1 but not F2 generations (Figure 2C). Note that early-generation *prg-1 daf-16* double mutants displayed wild-type levels of longevity (Figure 2), in contrast to *daf-16* single mutants which are known to be modestly short-lived (Kenyon et al. 1993).

Loss of germ cells through laser ablation or mutation of *glp-1* results in sterile worms that are long-lived (Berman & Kenyon 2006). We asked if *prg-1* mutant longevity might be due to the *glp-1* ‘germline aging’ pathway. Sterile *glp-1* mutants raised at the restrictive temperature of 25°C and fertile late-generation *prg-1* mutants displayed significant increases in lifespan when compared to wildtype (Figure 2D, p<0.0001). When fertile late-generation *prg-1; glp-1* double mutants were raised at the restrictive temperature of 25°C to induce sterility, they displayed a significant increase in lifespan when compared to late-generation *prg-1* single mutant controls (p=0.0001). However, this increase in lifespan was not additive and suggests that the longevity of late-generation *prg-1* mutants with strong reductions in fertility partially overlaps with the germline aging pathway.

To confirm the above results, we examined the longevity of *prg-1* mutants that are deficient for the *daf-12* nuclear hormone receptor, which is essential for the germline aging pathway (Hsin & Kenyon 1999). *daf-12* mutation significantly decreased late generation *prg-1* mutant lifespan (Figure 2E, p<0.05), but late generation *prg-1; daf-12* double mutants were significantly longer lived than *daf-12* single mutants (Figure 2E, p<0.001). These results confirm that late-generation *prg-1* mutants live long via a pathway that partially overlaps with but is distinct from the germline aging pathway.

To further explore the role of insulin signaling in *prg-1* mutants, we created *prg-1; daf-16::gfp* mutant lines using a *daf-16::gfp* transgene (Henderson & Johnson 2001). We found that early-generation adults displayed intestinal DAF-16::GFP expression that was primarily cytoplasmic, similar to *daf-16::gfp* transgene controls (Figure 3A,D) (Lin et al., 2001). Late-generation *prg-1* mutants that live long displayed mildly nuclear DAF-16::GFP localization in somatic cells (Figure 3B,D), whereas sterile *prg-1* mutant adults displayed strong nuclear localization of DAF-16::GFP in somatic cells, similar to that observed with *daf-16::gfp; daf-2* mutant controls, where DAF-16 is constitutively activated (Figure 3C,D).

**Figure 3.**
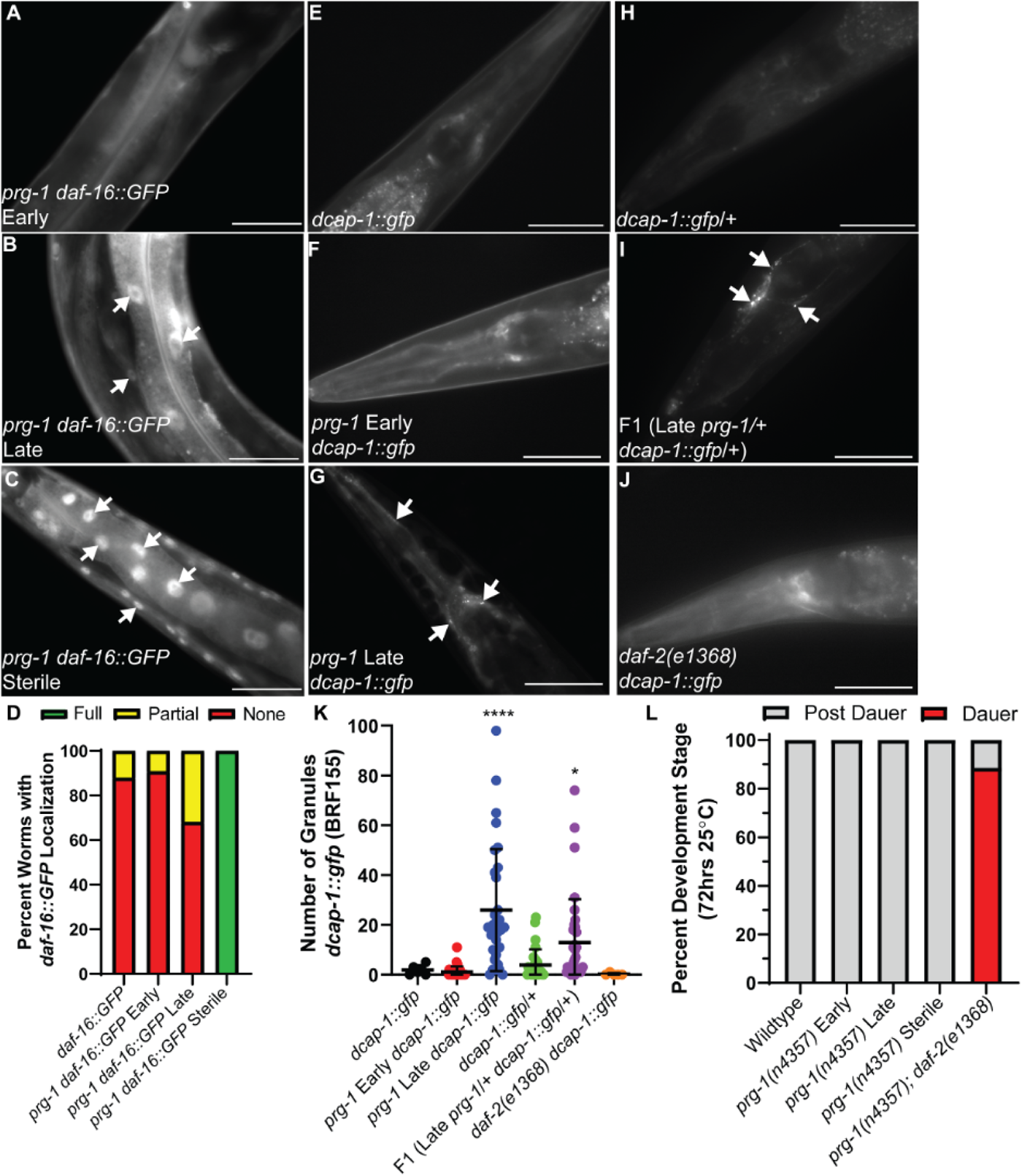
Induction of somatic stress response with heritable stress. Intestinal and muscular DAF-16::GFP remains cytoplasmic in early generation *prg-1(n4357)* mutants (A), but demonstrates partial nuclear localization in late generation animals (B) and strong or full nuclear localization in sterile generation *prg-1* mutant adults (C). White arrows indicate nuclear localization. (D) Percent of wildtype, early generation *prg-1*, late generation *prg-1*, and sterile generation *prg-1* worms with no, partial, or full *daf-16*::GFP nuclear localization in somatic tissue. (n=25, 22, 22, 20 respectively). Representative images of P-body granules visualized using *dcap-1::gfp* in wildtype (E), early *prg-1(n4357)* (F), late generation *prg-1(n4357)* (G), F1 generation of *dcap-1::gfp* males crossed to wildtype (H), F1 generation of *dcap-1::gfp* males crossed to late generation *prg-1(n4357)* (I), and *daf-2(e1368)* (J). White arrows indicate granules (K) Number of P-body granules in the head of worms represented in (E-J). (L) Percent worms of each genotype at either dauer or larval stages after dauer after 72 hours at 25°C. Scale bars = 50 µm. * p<0.05, **** p<0.0001.

### Transgenerational induction of P-bodies but not dauer larvae in *prg-1* mutants

Processing bodies (P-bodies) and stress granules are cytoplasmic aggregates composed of proteins and RNA that regulate mRNA translation and turnover (Rousakis et al. 2014). P-bodies appear under stressful conditions, including forms of environmental stress such as heat shock, radiation, osmotic, oxidative and ER stress (Protter & Parker 2016). Transposons that are transcriptionally suppressed by PRG-1 are parasites that resemble viruses, which can evoke strong host stress responses. Given that transposons are expressed in long-lived late-generation *prg-1* mutants (Simon et al. 2014), and given our hypothesis that *prg-1* mutants may transmit a heritable stress (Heestand et al. 2018), we monitored P-body abundance by crossing a *dcap-1::gfp* transgene (Rousakis et al. 2014) into *prg-1* mutant backgrounds and assessing DCAP-1::GFP granule expression in early- and late-generation strains. Independent *prg-1; dcap-1::gfp* lines revealed that DCAP-1::GFP granules were rarely present in wildtype and early-generation *prg-1* mutant strains but were significantly induced in late-generation *prg-1* mutants with strongly reduced fertility (Figure 3 E-G,K, Figure S1 A-C,E). Additionally, when males containing a *dcap-1::gfp* transgene were crossed with late-generation *prg-1* mutants, their F1 cross progeny exhibited significant increases in P-body formation in comparison with controls (Figure 3H,I,K).

Given that the longevity of late-generation *prg-1* mutants with compromised fertility depends on DAF-16, we asked if P bodies are induced in response to DAF-16 activity using a *daf-2* mutation that constitutively activates DAF-16. In a *daf-2* mutant background, there was no induction of DCAP-1::GFP granules (Figure 3 J,K Figure S1 D,E). The protein composition of stress granules is distinct from that of P bodies (Protter & Parker 2016), so we asked if stress granules are induced in late generation *prg-1* mutants or *daf-2* mutants using the stress granule marker *gfp::tiar-1* but found that stress granules were not induced (Figure S1 F-J).

Environmental stresses such as temperature, crowding, and starvation can induce an alternative *C. elegans* developmental stage termed the dauer larva, which is a state of developmental arrest that is long-lived and highly stress resistant (Baugh & Hu 2020). As the induction of dauer larvae is an intrinsically temperature-sensitive process, we asked whether late-generation *prg-1* mutants give rise to dauer larvae at the stressful temperature of 25°C. We did not observe any dauer larvae in early-, late-, or sterile generation *prg-1* mutants at 25°C. However, many dauer larvae were induced in a *prg-1; daf-2* double mutant control strain (Fi.g 3L), which indicates that mutation of *prg-1* does not suppress dauer formation. The lack of dauer larvae in late-generation *prg-1* mutants suggests that reduced DAF-2 insulin/IGF-1 signaling is not responsible for their longevity.

### Knockdown of *dcap-1* shortens late-generation *prg-1* mutant lifespan

Given the accumulation of P-bodies of late generation *prg-1* mutants, we asked if their longevity was dependent on *dcap-1*. As a control, we tested *daf-16* RNAi and observed a significant shortening of lifespan in middle- and late-generation *prg-1* mutant strains in comparison to wild-type controls (Figure 4B, p<0.0001). Furthermore, *daf-16* RNAi resulted in late-generation *prg-1* mutants that were significantly shorter-lived than middle-generation *prg-1* mutants (Figure 4B, p<0.01). Similarly, middle- and late-generation *prg-1* mutants fed *dcap-1* RNAi showed significant shortening of lifespan compared to controls (Figure 4C, p<0.0001) and late-generation *prg-1* mutant animals were shorter lived than middle-generation animals (Figure 4C, p<0.001). However, middle- and late-generation *prg-1* mutants were not short-lived when subjected to RNAi targeting *tiar-1*, which encodes a stress granule component (Figure 4D).

**Figure 4.**
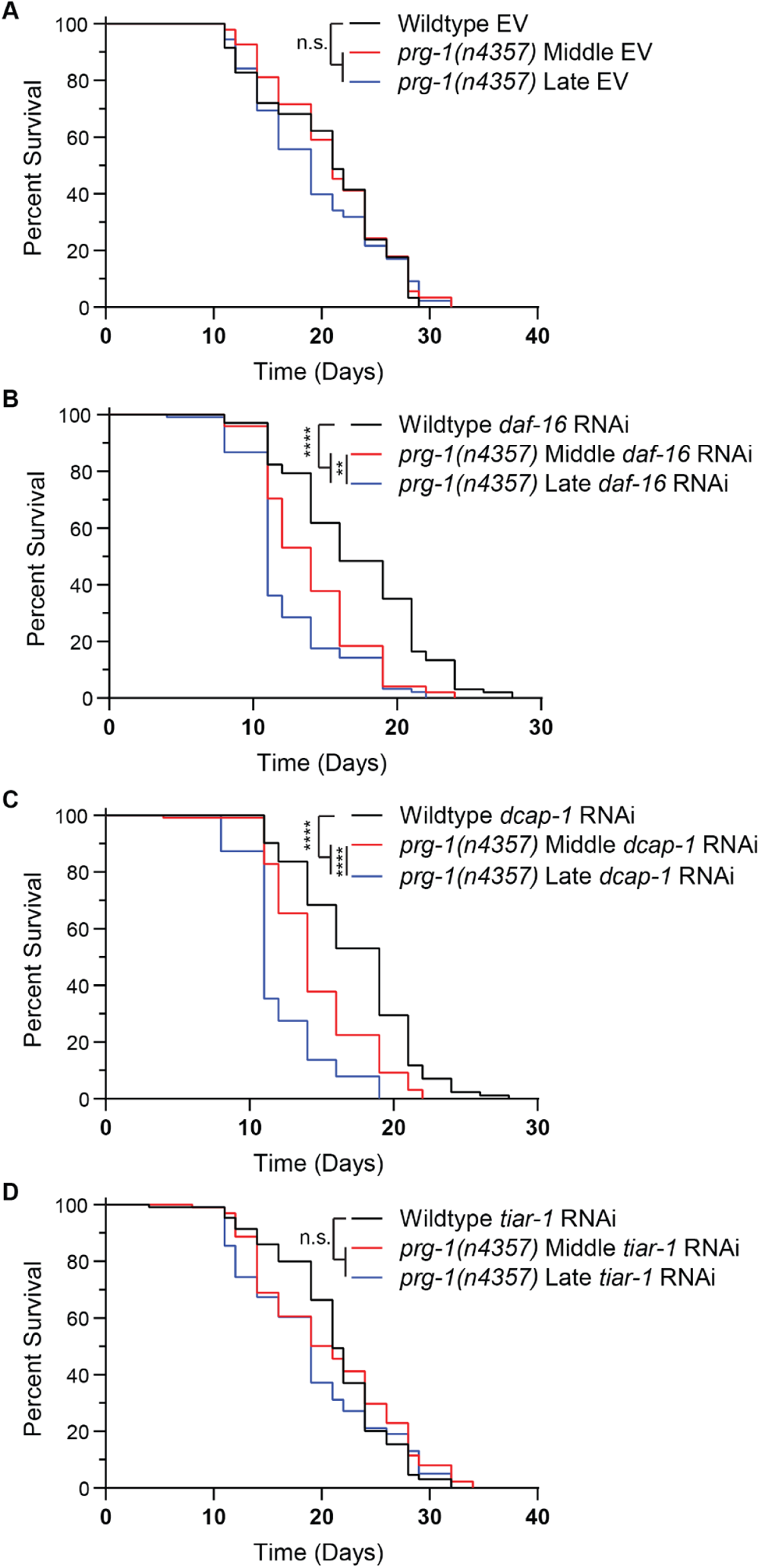
DAF-16 and DCAP-1 repress deleterious effects of *prg-1* mutant heritable stress. Lifespan curves for wildtype, middle generation *prg-1(n4357)*, and late generation *prg-1(n4357)* on (A) Empty Vector (EV) RNAi, (B) *daf-16* RNAi, (C) *dcap-1* RNAi, and (D) *tiar-1* RNAi. n.s. is not significant, **p<0.01, ****p<0.0001.

In our RNAi experiments, late-generation *prg-1* mutant worms fed empty vector RNAi (L4440) did not display an extension of lifespan compared to middle-generation and wild-type controls (Figure 4A). This was also true for late-generation *prg-1* mutants fed RNAi targeting *tiar-1* (Figure 4D). The lack of longevity of late-generation *prg-1* mutants grown on RNAi plates could be related to the thin crusty lawns of HT115 RNAi bacteria whose nutritional value is likely quite different from that of creamy, sticky lawns created by OP50 bacteria, which is the standard *C. elegans* food source used in most of our experiments (Revtovich et al. 2019). The shortened lifespan of late-generation *prg-1* mutants fed *daf-16* or *dcap-1* RNAi is consistent with our model that an endogenous heritable stress promotes the longevity of late-generation *prg-1* mutants, and that this stress becomes harmful in the absence of DAF-16 or DCAP-1 stress response factors.

### Long-lived late-generation *prg-1* mutants have small germline nucleoli

Small somatic cell nucleoli are a hallmark of longevity and are required for the longevity of all tested long-lived *C. elegans* mutants (Tiku et al. 2017). The size of the nucleolus reflects the level of rDNA transcription and the epigenetic state of a significant fraction of the nucleus (Tiku et al. 2017). Altered nucleolar size can reflect an epigenetic change that concerns rRNA synthesis and ribosome biogenesis. Moreover, the fertility defect of late-generation *prg-1* mutants is associated with increased levels of small RNAs antisense to rRNA and with altered rDNA copy number (Wahba et al. 2021). We therefore examined the nucleolar size of *prg-1* mutant germ cells. As somatic cells of *daf-2* mutants have small nucleoli (Tiku et al. 2017), we initially tested an allele of *daf-2, e1368,* which is commonly used for studies of aging (Kenyon et al. 1993). *daf-2(e1368)* mutants were long-lived, and their longevity was significantly greater than that of late-generation *prg-1(tm872)* mutants (Figure 5C, p<0.0001). However, *daf-2(e1368)* mutants had germ cell nucleoli that were normal in size (Figure 5A). In contrast, analysis of independent *prg-1* mutations revealed late-generation animals with small germline nucleoli in comparison to early-generation *prg-1* mutants and wild-type controls (Figure 5A,B). Furthermore, small germline nucleoli were transmitted to the F1 cross progeny but not to the F2 progeny of long-lived late-generation *prg-1* mutants (Figure 5A,B). Although we did not ask if somatic nucleoli of long-lived late-generation *prg-1* mutants and their F1 cross-progeny are small, we anticipate that this is the case because every known long-lived *C. elegans* mutant tested possesses small somatic nucleoli (Tiku et al. 2017).

**Figure 5.**
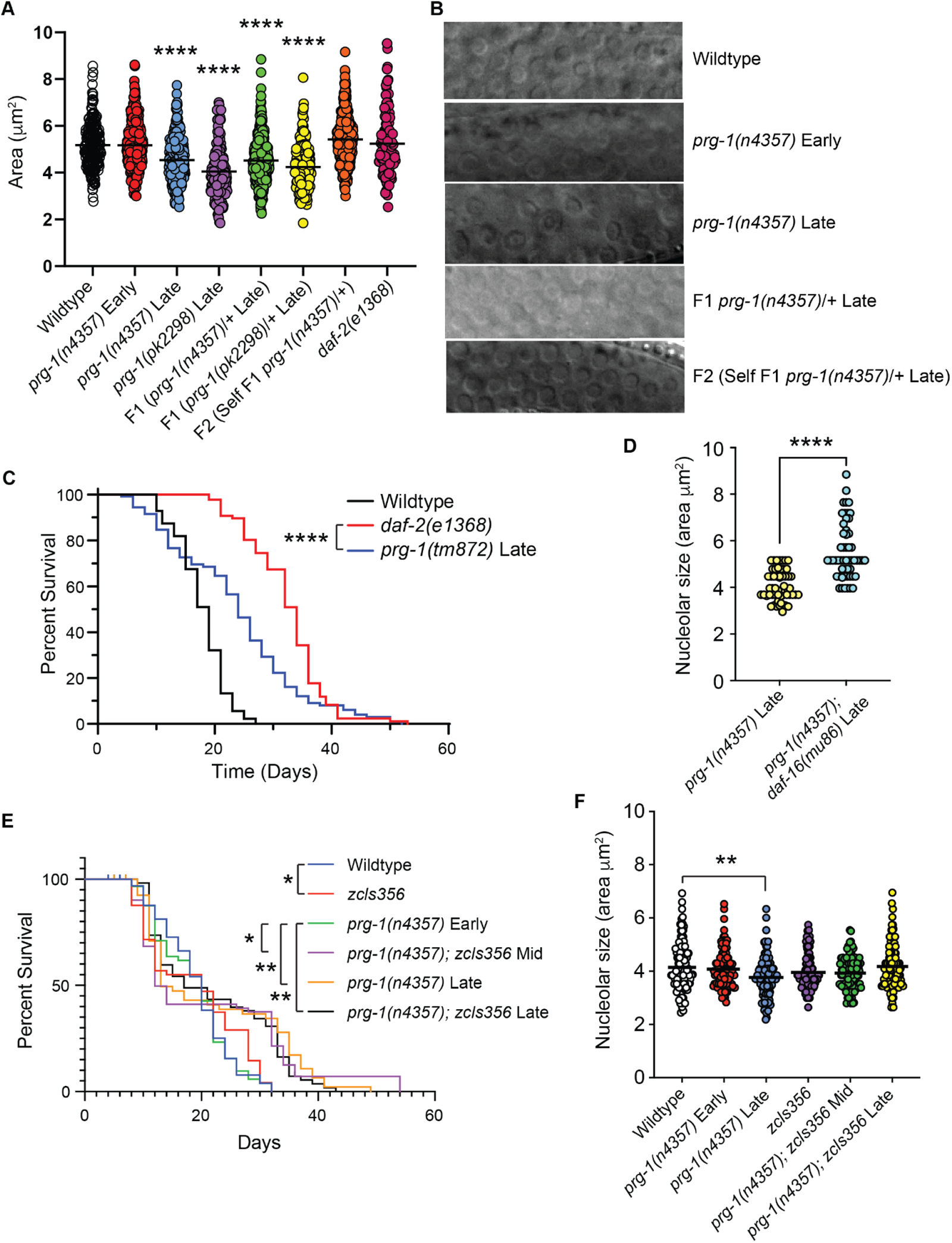
Small germline nucleoli of late-generation *prg-1* mutants. (A) Area measurements of mitotic germline nucleoli at the distal tip. (B) Representative images of germline nucleoli. (C) Lifespan curve for wildtype, *daf-2(e1368)*, and late generation *prg-1(tm872)*. ****p<0.0001. (D) Germline silencing of *daf-16* with *zcIs356* transgene abolishes small nucleoli of late-generation *prg-1* mutants. (E) Mid- and late-generation *prg-1; zcIs356* adults live long.

**Figure 6.**
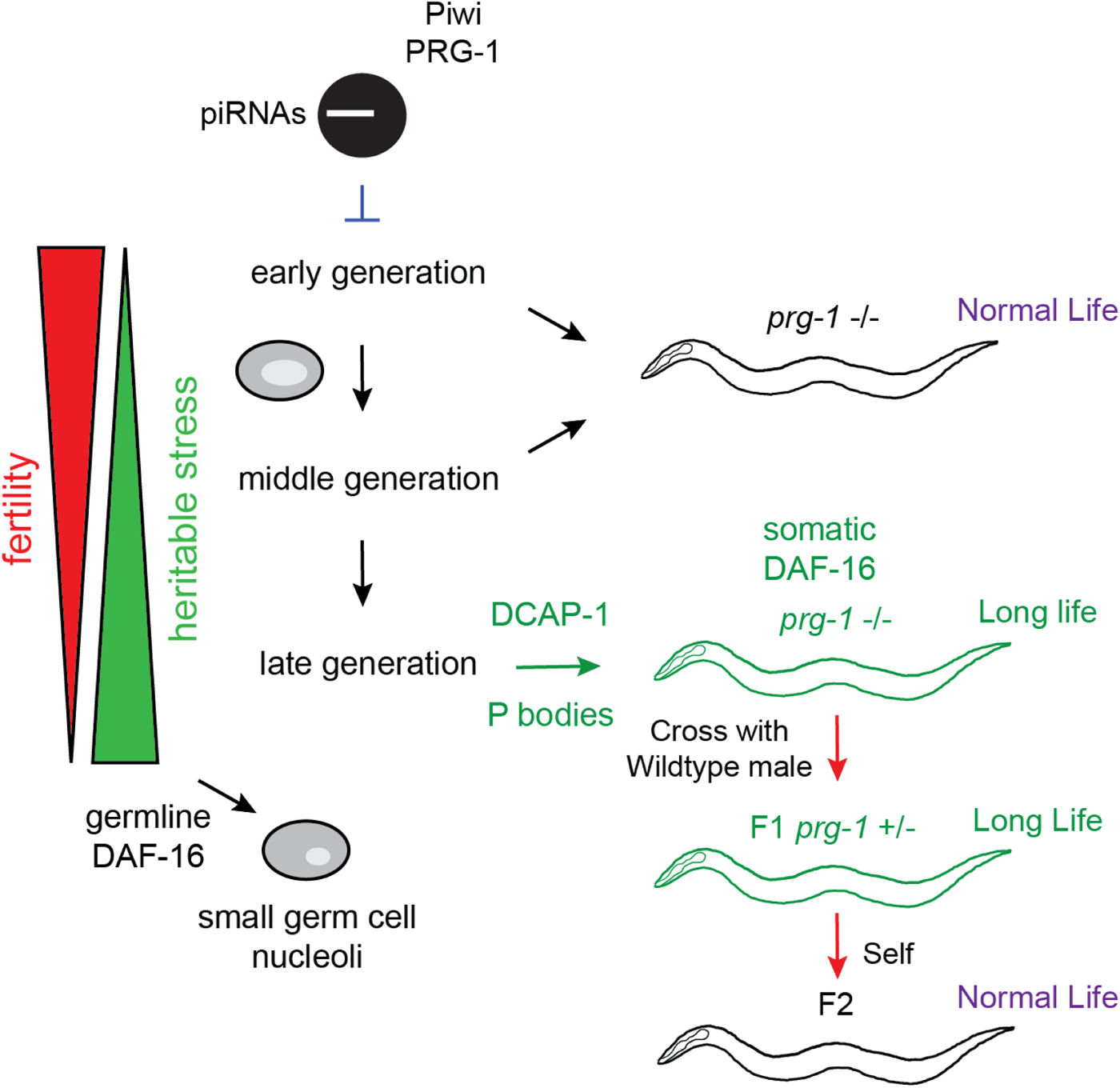
Effects of heritable stress transmitted by *prg-1* mutant germ cells. Loss of *prg-1* leads to a transgenerational reduction in fertility and increased levels of heritable stress (left). In late-generation *prg-1* mutant animals, high levels of heritable stress promote somatic P-body formation through DCAP-1, and both DCAP-1 and DAF-16/Foxo promote longevity and repress toxic effects of heritable stress in late-generation *prg-1* mutants. Small germ cell nucleoli represent an epigenetic response to the heritable stress transmitted by *prg-1* mutant germ cells.

Given that DAF-16 promotes *prg-1* mutant longevity, we examined germ cell nucleoli of short-lived late-generation *prg-1; daf-16* double mutants and observed that they were not reduced in size (Fig. 5D). We asked if DAF-16 acts in germ cells to promote small germ cell nucleoli of late-generation *prg-1* mutants using an integrated high copy number transgene, *zIs356*, which expresses high levels of DAF-16::GFP in somatic cells and rescues longevity when *daf-16* is deficient (Henderson & Johnson 2001). High copy number transgenes like *zIs356* can be silenced in germ cells (Ketting & Plasterk 2000; Dernburg et al. 2000). Because *zIs356 adults* express high levels of background somatic DAF-16::GFP fluorescence, we separated germlines of *zIs356* adults from somatic cells by microdissection. Although we did not observe DAF-16::GFP expression in oocytes of *zIs356* germlines, control *ot971* animals that have GFP inserted in the endogenous *daf-16* locus displayed DAF-16::GFP in oocytes (Figure S2 A-D) (Aghayeva et al. 2020). We hypothesized that lack of DAF-16::GFP fluorescence in *zIs356* germ cells might be a consequence of silencing of repetitive transgenes in germ cells via cosuppression, where small RNAs silence both the transgene and the endogenous locus (Ketting & Plasterk 2000; Dernburg et al. 2000). To test this hypothesis, we crossed *ot971* males that express endogenous *daf-16::GFP* with *zIs356* hermaphrodites and observed that their F1 cross-progeny did not express DAF-16::GFP in oocytes (Figure S2 E,F). We conclude the *zIs356* high copy number *daf-16::GFP* transgene promotes cosuppression-mediated silencing of both *zIs356* and the endogenous *daf-16* locus in germ cells.

We created independent *prg-1(n4357); zIs356* strains and propagated them for multiple generations to ask if germline DAF-16 affects either nucleolar size of germ cells or somatic longevity. We propagated *prg-1(n4357); zIs356* strains and observed a moderate reduction in brood size even in the earliest generations, even though *zIs356* controls displayed normal fertility. It was previously demonstrated that deficiency for *daf-16* enhances the fertility defect of *prg-1* mutants (Simon et al. 2014), and loss of *daf-16* in germ cells may synergize with overexpression of somatic DAF-16 to perturb *prg-1* mutant fertility. We therefore studied mid-generation *prg-1(n4357); zIs356* adults with moderately reduced fertility and late-generation *prg-1(n4357); zIs356* adults with strongly reduced fertility that were not sterile. *zIs356* adults and mid-generation *prg-1(n4357); zIs356* adults were modestly long-lived in comparison with wild-type or early-generation *prg-1* mutant controls, likely due to high levels of somatic DAF-16::GFP expression (Figure 5F). However, both late-generation *prg-1* mutants and late-generation *prg-1(n4357); zIs356* adults were long-lived.

Although nucleolar size of mitotic germ cells was reduced in late-generation *prg-1* mutant, nucleolar size of mid- and late-generation *prg-1(n4357); zIs356* strains was not different than that of *zIs356* controls (Figure 5E). Therefore, germline DAF-16 reduces the volume of germ cell nucleoli in late-generation *prg-1* mutants in a manner that is distinct from its role in somatic longevity. This represents one of few known roles for DAF-16 in germ cells.

The ratio of nucleolar to nuclear size has not been studied in the context of *C. elegans* adult longevity. We found that nuclear size of mitotic germ cells was wild-type for all *prg-1* and *prg-1(n4357); zIs356* strains, except for late-generation *prg-1(n4357); zIs356* adults that had modestly enlarged nuclei (Figure S3A). The ratio of nucleolar to nuclear size was modestly reduced for *zcIs356* and mid-generation *prg-1(n4357); zIs356* strains, and reduced early- and late-generation *prg-1* mutants as well as late-generation *prg-1(n4357); zIs356* strains that were long-lived (Figure S3B). Long-lived late-generation *prg-1* mutants had the smallest nucleolar to nuclear size ratio of any strain tested (Figure S3B). We conclude that the ratio of nucleolar to nuclear size for mitotic germ cells correlates with adult longevity. However, this relationship is more complex than that of nucleolar size alone.

## Discussion

The disposable soma theory of aging posits that trade-offs occur when allocating resources to reproduction or somatic longevity (Kirkwood & Holliday 1979). We previously found that sterile late-generation *prg-1* mutant adults are in a state of reproductive arrest (Heestand et al. 2018). Adult reproductive arrest can occur in response to the stress of acute starvation and is accompanied by adult longevity (Angelo & Van Gilst 2009). We therefore studied *prg-1* mutants longevity and discovered that both fertile and sterile late-generation *prg-1* mutants were long-lived. In contrast, we previously observed normal longevity for fertile and sterile late-generation *C. elegans trt-1* telomerase mutants (Meier et al. 2006). Therefore, the reduced fertility of fertile late-generation *prg-1* mutants, *per se*, does not explain their longevity.

Although late-generation *prg-1* mutants display reduced fertility, experiments with *glp-1(e2141)* and the *daf-12* nuclear hormone receptor (Figure 2 D,E) revealed that longevity of late-generation *prg-1* mutants is distinct from pathway that promotes longevity in response to germline ablation (Hsin & Kenyon 1999). We instead found that loss of the DAF-16/Foxo transcription factor caused late-generation *prg-1* mutants to live shorter lives than short-lived *daf-16* single mutants (Kenyon et al. 1993; Baugh & Hu 2020). Loss of *daf-16* caused mid-generation *prg-1* mutants with mild fertility defects to live very short lives, and this also occurred when the P body factor *dcap-1* was knocked down. Given that P bodies and DAF-16 are upregulated in response to stress, the longevity of late-generation *prg-1* single mutants resembles a hormetic stress response (Calabrese et al. 2015; Shore & Ruvkun 2013; Masoro 2007). We propose that the longevity of late-generation *prg-1* mutants is due to heritable stress, an unidentified epigenetic factor that is transmitted by *prg-1* mutant germ cells and accumulates over generations to reduce fertility and promote longevity. High levels of heritable stress present in late-generation *prg-1* mutants that display strong reductions in fertility induce somatic longevity (Figure 1). Moderate levels of heritable stress that are normally inconsequential for longevity become debilitating and markedly shorten lifespan for mid-generation *prg-1* mutant animals in the absence of DAF-16 or DCAP-1 stress response factors (Figures 2A, 4B,C). Moderate levels of heritable stress are sufficient to trigger longevity in mid-generation *prg-1(n4357); zIs356* adults that overexpress somatic DAF-16 (Figure 5E). This speaks to an interplay between endogenous stress response pathways and levels of heritable stress transmitted by germ cells that might be generally relevant to the epigenetic regulation of aging in metazoans.

Although the DAF-16 transcription factor is required *daf-2 i*nsulin/IGF-1 mutant longevity, late-generation *prg-1* mutants did not display dauer larvae that are commonly produced under circumstances of stress at the stressful temperature of 25°C, which strongly promotes dauer formation for *daf-2* mutants (Figure 3L) (Baugh & Hu 2020). Moreover, *daf-2* mutants that are deficient for *daf-16* do not live shorter than *daf-16* single mutants (Kenyon et al. 1993), in contrast to very short-lived late-generation *prg-1; daf-16* double mutants. Together, these results indicate that reduced *daf-2* insulin/IGF-1 signaling is not responsible for the longevity of late-generation *prg-1* mutants.

*prg-1* mutant germ cells transmit a form of stress that stochastically accumulates over generations to cause reduced fertility and ultimately reproductive arrest (Simon et al. 2014; Heestand et al. 2018). We crossed late-generation *prg-1* mutants with strong reductions in fertility with wild-type males and found that F1 but not F2 cross progeny of late-generation *prg-1* mutants were long-lived (Figure 1), indicating that the heritable epigenetic factor transmitted by *prg-1* mutant gametes promotes longevity for a single generation if wild-type PRG-1/Piwi is restored. Inactivation of members of the ASH-2 trithorax complex that promotes histone H3K4 methylation elicits an adult longevity phenotype that is transmitted for three generations to genetically wildtype progeny (Greer et al. 2011). In addition, ASH-2 trithorax inactivation extends *daf-16* mutant longevity (Greer et al. 2016), whereas late-generation *prg-1* mutants are very short-lived in the absence of *daf-16* (Figure 2A, 4B). Therefore, the heritable longevity observed in late-generation *prg-1* mutants and in response to ASH-2/trithorax deficiency occur in response to distinct epigenetic mechanisms.

The SPR-5 H3K4 demethylase is required for germ cell immortality, and late-but not early-generation *spr-5* mutants are long-lived (Greer et al. 2016). Late-generation *spr-5* mutant longevity was partially additive with longevity induced by *glp-1* germline ablation, as observed for late-generation *prg-1* mutants (Figure 2D). However, *daf-12* was completely required for late-generation *spr-5* mutant longevity (Greer et al. 2016), whereas *daf-12* is only partially required for the longevity of late-generation *prg-1* mutants (Figure 2E). Moreover, *daf-16* is not required for the longevity of late-generation *spr-5* mutants (Greer et al. 2016). This implies that deficiency for *prg-1* and *spr-5* result in transgenerational accumulation of distinct epigenetic factors that promote longevity.

Several causes of sterility for late-generation *prg-1* mutants have been proposed, including misrouting of pro- and anti-silencing small RNAs (de Albuquerque et al. 2015), expression of high levels of transposon or repetitive RNAs (Simon et al. 2014), transposition of the Mirage transposon (McMurchy et al. 2017), disruption of gene expression (Reed et al. 2020), excessive silencing of histone genes (Barucci et al. 2020) and disruption of germ granules (Spichal et al. 2021). A recent intriguing study suggested that the fertility of late-generation *prg-1* mutants may be compromised by accumulation of high levels of small RNAs that are homologous to rRNA as well as the presence of extrachromosomal rDNA circles (Wahba et al. 2021), although rRNA levels appear unaffected in *prg-1* mutants (Reed et al. 2020). We discovered that germ cells of late-generation *prg-1* mutants and their F1 cross-progeny have small nucleoli that are created in response to germline DAF-16 activity but that do not impact *prg-1* mutant longevity. The small mitotic germ cell nucleoli of late-generation *prg-1* mutants instead reflect a DAF-16-mediated epigenetic response to the heritable stress transmitted by late-generation *prg-1* mutant germ cells. *Drosophila* Piwi becomes localized to the nucleolus in response to heat stress, suggesting a potentially conserved relationship between Piwi/PRG-1 and rDNA (Mikhaleva et al. 2019).

We defined three traits found in late-generation *prg-1* mutants and in their F1 but not F2 cross progeny: reduced fertility, increased levels of DCAP-1 P-body foci, and reduced germline nucleoli (Figures 3 and 5A,D). *C. elegans* P-bodies increase in aged wild-type animals, in response to oxidative stress, heat shock, starvation, hypoxia, and in response to deficiency for the XRN-1 mRNA exonuclease that results in global impairment of RNA degradation (Rousakis et al. 2014). Deficiency for *dcap-1* substantially compromises survival of wildtype in response to oxidative stress or heat shock (Rousakis et al. 2014). In contrast, P-bodies are strongly repressed in somatic cells of long-lived *ife-2* mutants (Rousakis et al. 2014), which are deficient for a translation initiation factor whose loss leads to reductions in global protein synthesis (Syntichaki et al. 2007). We similarly found that P-body levels were not increased in a *daf-2* mutant background where longevity is dependent on DAF-16, which suggests that high P-body levels of late-generation *prg-1* mutants are not a general consequence of upregulated DAF-16 activity (Figure 4A-C). The low P-body levels present in long-lived mutants like *daf-2* and *ife-2* could reflect low levels of endogenous stress.

In contrast to long-lived *ife-2* or *daf-2* mutants, strong increases in P-body formation occur in long-lived late-generation *prg-1* mutants and their F1 cross progeny, and deficiency for either *daf-16* or *dcap-1* caused late-generation *prg-1* mutants to be substantially shorter lived than short-lived controls that lack *daf-16* or *dcap-1* (Figure 3, Figure S1). In contrast, *dcap-1* deficiency only modestly shortens the lives of most if not all long-lived *C. elegans* mutants including *ife-2*, *daf-2*, *rsks-1* (TOR deficient) and *glp-1* (germline deficient) (Rousakis et al. 2014). This implies that the longevity of late-generation *prg-1* mutant adults is distinct from that of most known longevity pathways. Given that P-body formation occurs in response to various environmental stresses, the high levels of P-bodies in late-generation *prg-1* mutants could result from an endogenous response to heritable stress that is hormetic in nature and reflects one of several changes to small RNA and RNA metabolism that occur when *prg-1* is deficient (Simon et al. 2014; Barucci et al. 2020; Reed et al. 2020; de Albuquerque et al. 2015; Wahba et al. 2021).

It was recently demonstrated that *daf-16; dcap-1* double mutants display short lifespans that are not shorter than the short lifespans of either single mutant, indicating that DAF-16 and DCAP-1/P-bodies promote wildtype longevity via a common pathway (Borbolis et al. 2020). Neuronal overexpression of the P-body protein DCAP-1 is sufficient to promote stress resistance and longevity in a manner that requires *daf-16* (Borbolis et al. 2020). Somatic P bodies may therefore promote resilience and longevity in response to heritable stress transmitted by stimulating somatic DAF-16/Foxo in late-generation *prg-1*/Piwi mutant germ cells.

P-bodies can promote viral resistance and aspects of the innate immune response (Protter & Parker 2016), which are also properties of perinuclear germ granules that house the PRG-1/Piwi/piRNA genome silencing proteins (Reed et al. 2020). The reproductive arrest phenotype (sterility) of late-generation *prg-1*/Piwi mutants is associated with and likely triggered by dissolution of perinuclear germ granules (Spichal et al. 2021). Given that germ granules are RNA processing bodies and that some germ granule proteins also occur in P bodies (Cassani & Seydoux 2022), it is possible that high levels of somatic P bodies observed in late-generation *prg-1* mutants may be linked to germ granule perturbations of late-generation *prg-1* mutants.

Inactivation of the Piwi/piRNA pathway in vertebrates and *Drosophila* leads to immediate sterility, where acute effects of this germline epigenetic silencing system on longevity might occur. Inactivation of *Drosophila* Piwi results in decreased longevity and in reduced proliferation of adult somatic stem cells (Jones et al. 2016; Sousa-Victor et al. 2017). The increased longevity of late-generation *C. elegans prg-1*/Piwi mutants could be related to the lack of somatic stem cells in *C. elegans* adults. In addition, *Drosphila* Piwi mutants are immediately sterile, whereas strains that lack *C. elegans prg-1*/Piwi need to grow for many generations before markedly reduced fertility and increased longevity are observed. The lack of immediate sterility in *C. elegans* Piwi mutants might be due to an exceptional memory of Piwi-mediated genome silencing marks in nematode germ cells. We note that the *C. elegans* heterochromatin mutant *set-32* that is deficient for H3K23me3 silencing mark was recently shown to be long-lived (Huang et al. 2022), which is in line with heterochromatin deficiency as a potential cause of *prg-1*/Piwi mutant longevity.

The transgenerational sterility of *prg-1* mutants suggests a defect that accumulates as germ cells proliferate over generations, which is potentially analogous to proliferative aging or senescence of mammalian somatic cells (Micco et al. 2021). Several links between aging and dysregulation of the epigenome have been documented, including desilencing of retrotransposons in senescent human cells and epigenetic silencing of retrotransposons by the anti-aging protein SIRT6 in the mouse (Simon et al. 2019). The induction of longevity and reproductive arrest in response to an heritable epigenetic factor transmitted by *prg-1* mutant germ cells suggests that an analogous form of non-Mendelian inheritance (heritable stress) that might be transmitted to affect the rate of aging, fertility or adult onset disorders in mammals.

## Experimental Procedures

### Strains

Unless noted otherwise, all strains were cultured at 20°C on Nematode Growth Medium (NGM) plates seeded with *E. coli* OP50. Strains used include: Bristol N2 wildtype, *unc-13(e450) I*, *prg-1(n4357) I, prg-1(pk2298) I, prg-1 (tm872) I, daf-16(mu86) I*, OH16024 *daf-16(ot971[daf-16::gfp]), daf-2(e1368) III*, *glp-1(e2141) III*, TJ356 *zIs356* [*daf-16p::daf-16a/b::*GFP + *rol-6(su10060*], BRF155 *Pdcap-1::dcap-1::gfp::dcap-1 3’ UTR*, BRF261 *Pdcap-1::dcap-1::gfp::dcap-1 3’ UTR*, BRF211 *gfp::tiar-1*.

### Fertility of *prg-1* mutants does not depend on the precise generation number

*prg-1* mutations were outcrossed versus an outcrossed stock of the tightly linked mutation *unc-13(e450)*, and freshly isolated homozygous F2 lines were established for analysis. Freshly isolated F2 through F4 lines that were never starved were designated ‘early-generation *prg-1* mutants’. Middle generation worms were passaged as 6 L1s from early generation isolates each week until a small drop in brood size was observed without the presence of XO males. These phenotypes are typically observed from F10-F14 under our growth conditions, but they can arise many generations later. Stressed late generation *prg-1* mutants remained fertile but had very low brood sizes, a >2% frequency of XO males, and a low frequency of daughters with a Glp phenotype (adults that lack embryos and possess a clear patch in their uterus). Typically *prg-1* mutants with low or very low brood sizes were observed from generations F14 to F30. Sterile *prg-1* mutants remained sterile and gave rise to no progeny and had a Glp phenotype. Sterile *prg-1* mutants are primarily found in progeny of late-generation *prg-1* mutant parents whose brood size is low. Sterile *prg-1* mutants were typically observed within a few generations of the onset of low or very low brood sizes. To create the linked *prg-1 daf-16* double mutant, *prg-1 dpy-24* and *unc-13 daf-16* double mutants were first created, then progeny of *prg-1 dpy-24 / unc-13 daf-16* heterozygotes that lost *unc-13* were identified. The resulting putative *prg-1 daf-16* recombinant chromosomes were made homozygous by selecting against *dpy-24* and PCR genotyped to verify the presence of *prg-1* and *daf-16* deletions.

### Lifespan analysis

Animals were singled from non-starved plates and transferred every two days until the end of their reproductive cycles. Animals were kept uncrowded and free of contamination throughout lifespan experiments and transferred to new plates once per week. All animals were maintained at 20°C. For RNAi experiments, worms were transferred to their respective RNAi condition plates at the L3 stage.

### Statistical analysis

Statistical analysis was performed using the R statistical computing environment (Team & Others 2017) and the survival R package (Therneau & Others 2015). For lifespan experiments, pairwise Log-rank (Mantel-Cox) tests were performed to detect significant differences between groups. For the P-bodies experiment, a Kruskal-Wallace test was performed to detect the presence of any significant differences within the dataset. Pairwise Mann Whitney U tests were performed as a post-hoc test to detect the presence of significant differences between specific groups. All reported p-values were adjusted using Bonferroni correction to control the familywise error rate. Progeny count, nucleoli size, and DCAP-1::GFP aggregate statistics were done in Graphpad Prism 8.0 using Brown-Forsythe and Welch ANOVA tests and correcting for multiple comparisons.

### Nucleoli measurements

Animals were staged at larval stage 4 (L4) and Z stack images of the distal tip region of the germline were taken using a DeltaVision microscope (Applied Precision) with a 60X objective. Nucleoli size was measured on Fiji using the freehand draw tool. Nucleoli measurements were combined from >10 worms with multiple measurements taken from each germline.

## Acknowledgements

We thank members of the Ahmed lab for critical reading of the manuscript and P. Syntichaki and R. Dowen for strains. Some strains were provided by the CGC, which is funded by the NIH Office of Research Infrastructure Programs (P40 OD010440). This study was supported by NIH grant RO1 GM083048 (S.A) and by NIH R01 GM135470 (S.A.) and by NIH R01 ES035777. The funders had no role in study design, data collection and analysis, decision to publish, or preparation of the manuscript.

## Competing interests

Authors declare no competing interests.

## Author contributions

B.H., M.S., E.L., and B.M. performed experiments. B.M. and S.F. analyzed data. B.H., E.L., B.M. and S.A. wrote the manuscript. S.A. supervised the experiments.

## Data availability

Raw data for this study are deposited at Dryad: https://datadryad.org/stash/share/Oe8rDegiBr-tmMx7OJCwkQs8IfVXLhOsABH3tYsm5jI

---

**Supplemental Table 1.**
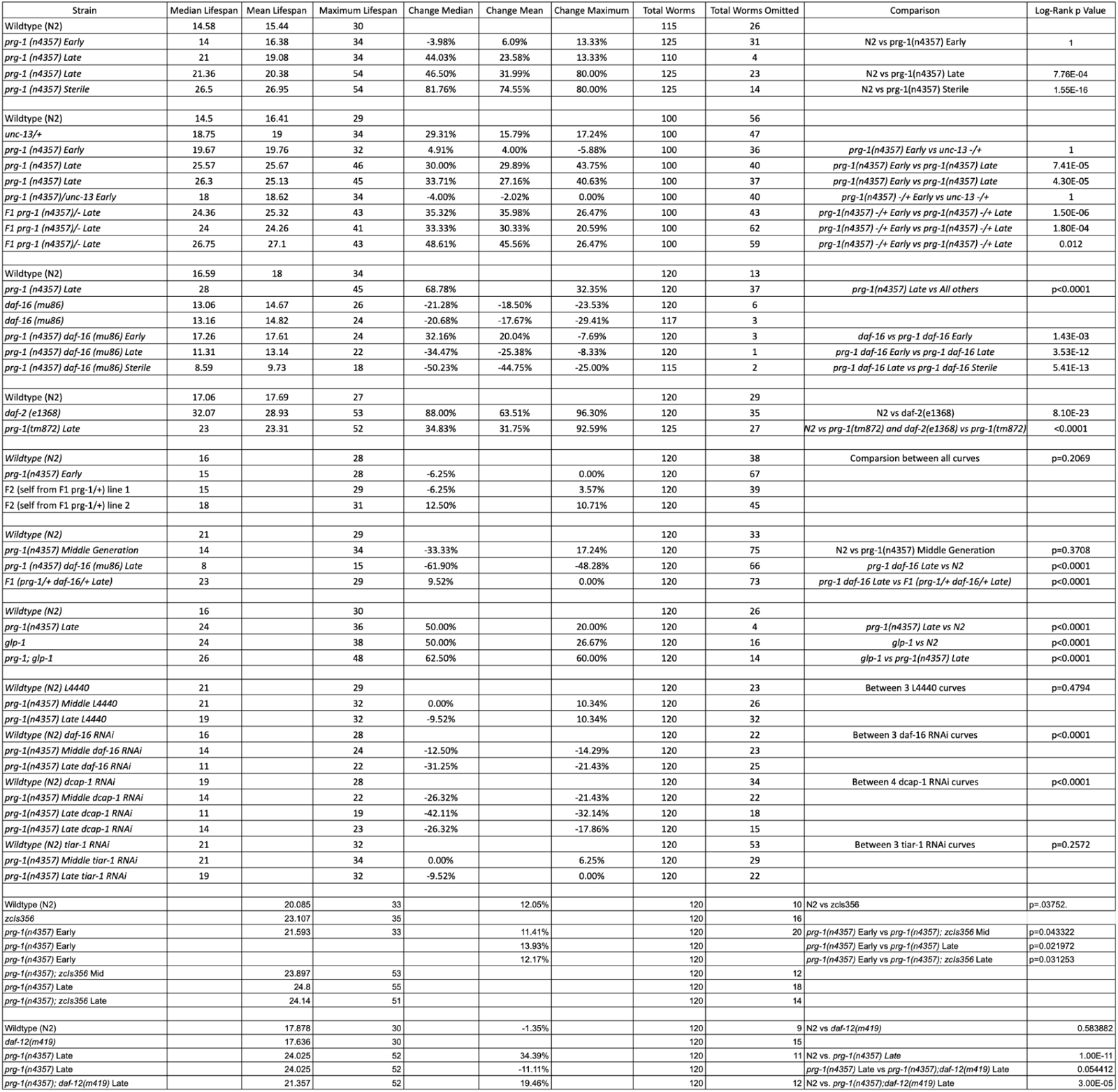
Analysis of aging assays. Log-rank (Mantel-Cox) analysis with Bonferroni correction comparing aging assays.

**Supplemental Figure S1.**
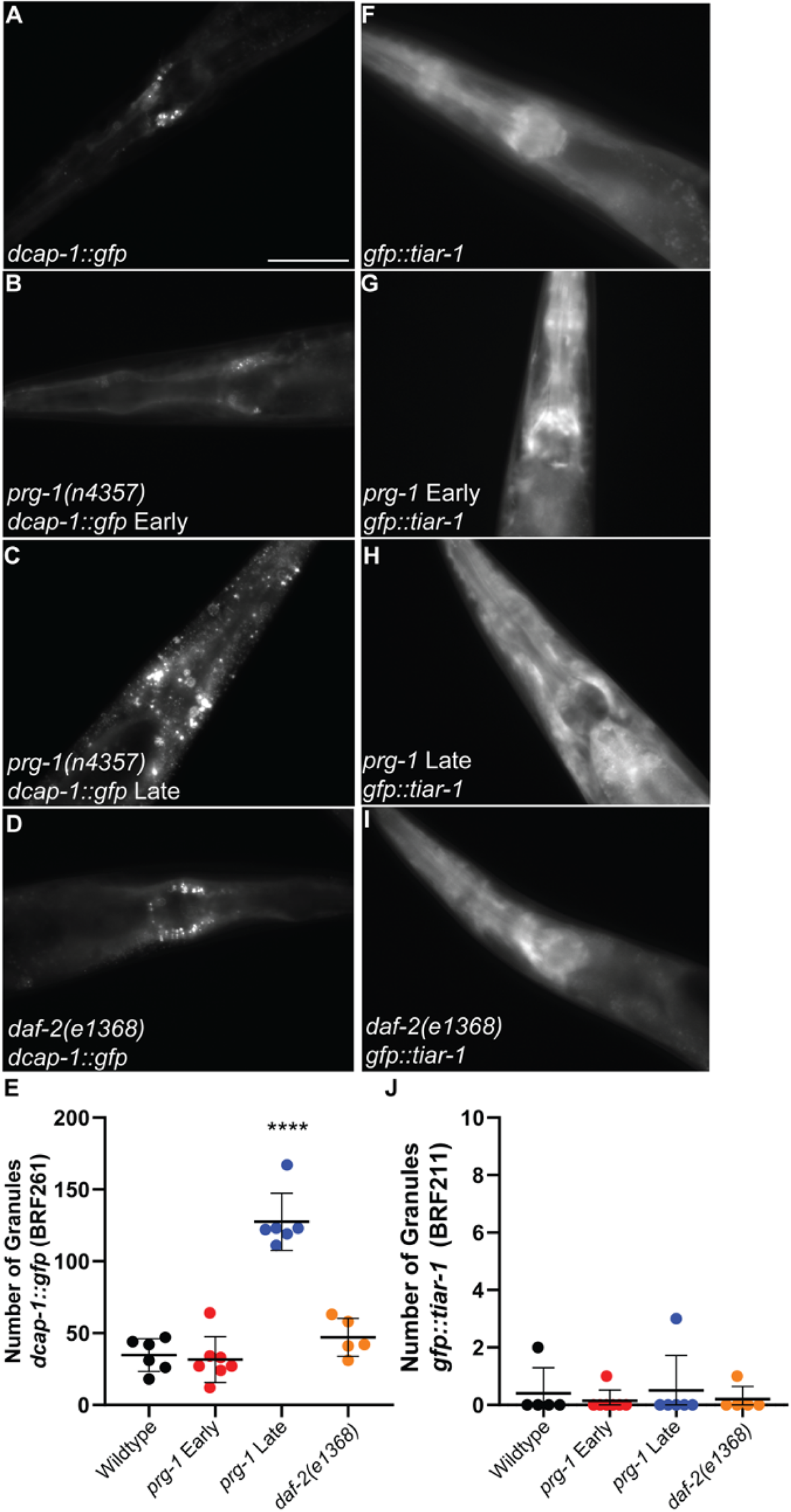
P-body granules are Induced in Late Generation *prg-1*. Representative images of P-body granules visualized using *dcap-1::gfp* (BRF261) in wildtype (A), early *prg-1(n4357)* (B), late generation *prg-1(n4357)* (C), and *daf-2(e1368)* (D). (E) Total number of granules in the head of worms represented in (A-D). Representative images of stress granules visualized using gfp::tiar-1 (BRF211) in wildtype (F), early *prg-1(n4357)* (G), late generation *prg-1(n4357)* (H), and *daf-2(e1368)* (I). (E) Total number of granules in the head of worms represented in (F-I). ****p<0.0001

**Supplemental Figure S2.**
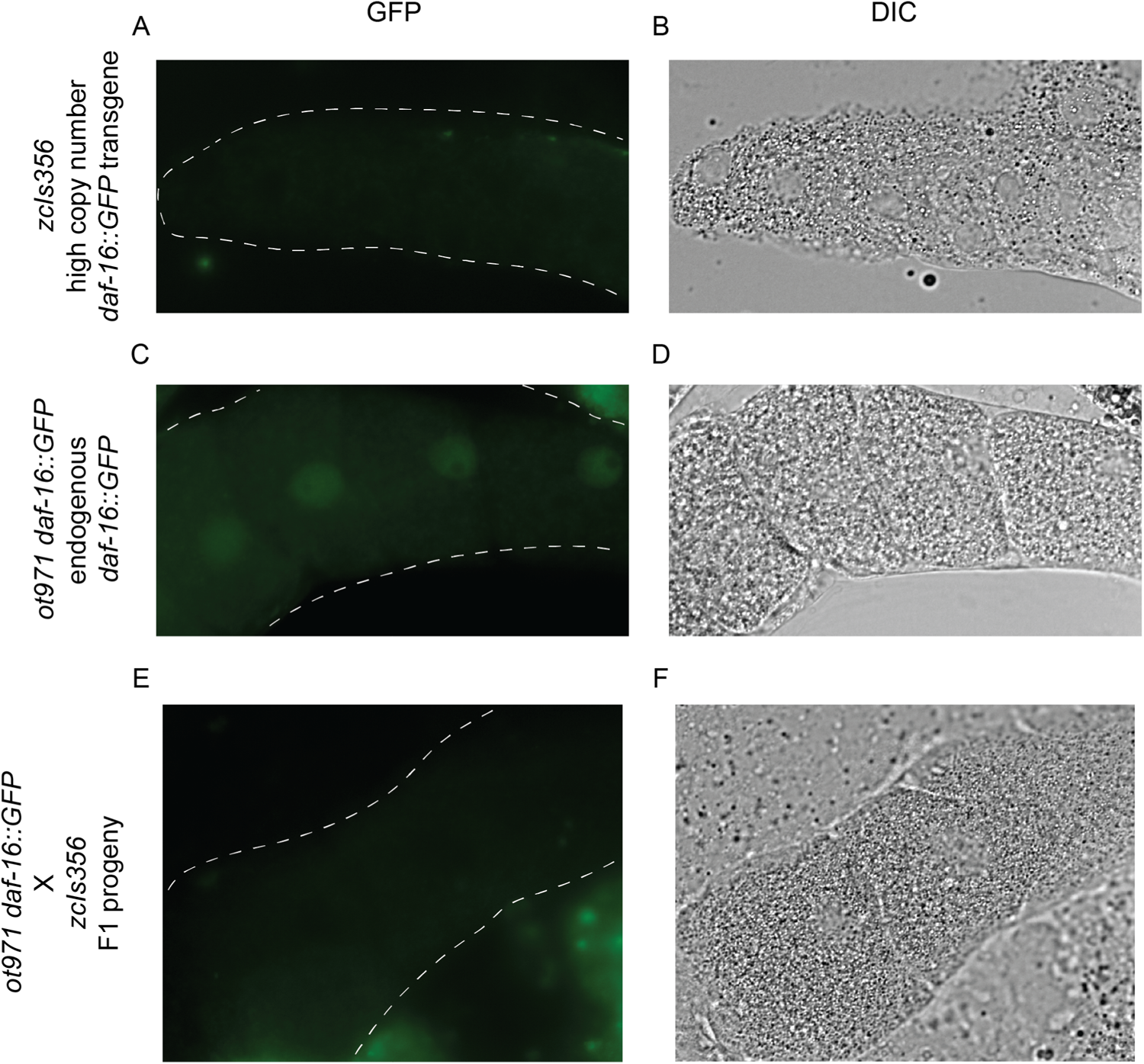
Silencing of *daf-16::GFP* in germ cells by high copy number *zcIs356 daf-16::GFP* transgene. (A-B) Oocytes of *zcIs356* adults fail to display GFP fluorescence. (C-D) Germ cells of *ot971* endogenous *daf-16::GFP* display faint nuclear DAF-16::GFP fluorescence. (E-F) F1 cross-progeny of *ot971* endogenous *daf-16::GFP* males crossed with a *zcIs356* hermaphrodite fail to display GFP in oocytes. Dashed white lines outline the location of oocytes in the GFP panels.

**Supplemental Figure S3.**
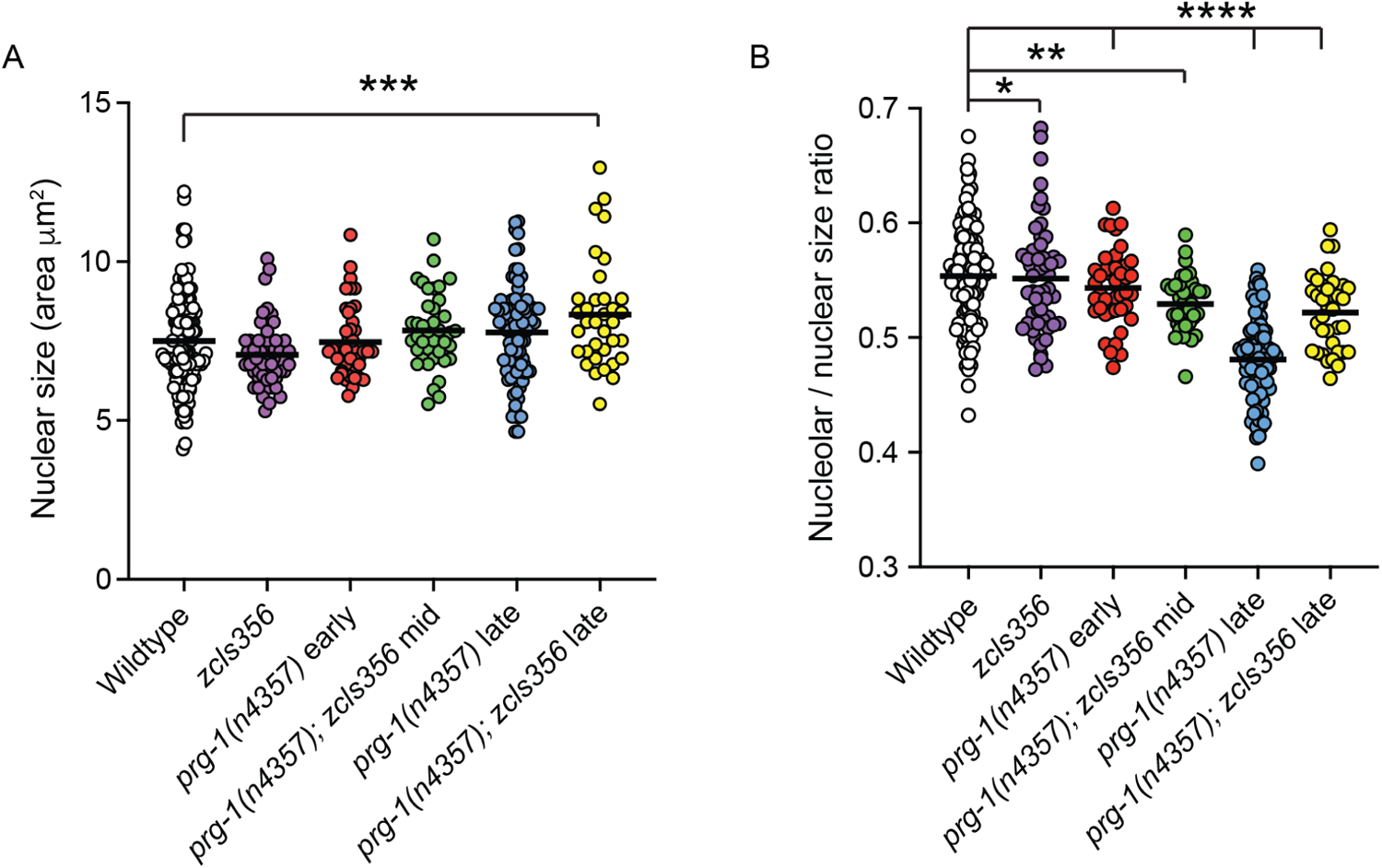
Analysis of nuclear / nucleolar size. (A) Nuclear size modestly increased in late-generation adult germ cells of a *prg-1; zcIs356* strain. (B) Nucleolar / nuclear size ratio is reduced in late-generation *prg-1* mutant germ cells.

## Notes

### Competing Interest Statement

The authors have declared no competing interest.

### Summary of Updates

Our progress includes confirmation that the germline aging pathway only partially overlaps with the pathway that promotes longevity in late-generation prg-1 mutants, based on a glp-1 mutation that activates the germline aging pathway and a daf-12 mutation that inactivates the germline aging pathway. We further demonstrate that small germline nucleoli of late-generation prg-1 mutants are created by a function of the DAF-16 longevity TF that is autonomous to germ cells. We demonstrate that the small germ cell nucleoli are not required for prg-1 mutant longevity, but are instead a response to the heritable stress transmitted by prg-1 mutants. This represents one of the few known functions of DAF-16 in germ cells.

